# Comparative Mapping of Functional and Structural Homologies in the Pig and Human Brain

**DOI:** 10.64898/2026.04.22.720235

**Authors:** Qiang Li, Ishfaque Ahmed, Moria F. Taber, Morgan H. Laballe, Sydney E. Sneed, Erin E. Kaiser, Franklin D. West, Qun Zhao, Vince D. Calhoun

## Abstract

Comparative mapping of functional and structural homologies across humans, small animals, and nonhuman primates has been extensively pursued due to its strong translational relevance. However, these experimental models possess inherent limitations in fully recapitulating the complexity of human cortical organization. The porcine model has recently emerged as a promising alternative, given its neuroanatomical and physiological similarities to the human brain. Despite these advantages, systematic cross-species characterization of functional and structural homologies between humans and pigs remains largely understudied. In the present study, we acquired resting-state functional MRI and diffusion MRI data from pigs and analyzed them alongside corresponding human datasets to investigate cross-species correspondence in large-scale brain organization. First, to enhance functional network alignment across species, group independent component analysis was performed separately within each species to identify intrinsic large-scale functional networks. Our results demonstrated that multiple canonical human resting-state networks are represented in the porcine brain, including sensorimotor, default mode, cerebellar, frontal, and central executive networks. Moreover, we observed significant cross-species concordance in intrinsic functional architecture across multiple distributed networks, both in spatial distribution and temporal patterns, indicating homologous large-scale brain organization between pigs and humans. Second, we conducted comparative structural analyses using diffusion MRI-based fiber tractography, color-encoded fractional anisotropy maps, and structural connectivity analyses to examine and compare white matter organization in pigs and humans. Cross-species comparison revealed substantial similarities in major white matter pathways and their spatial organization, supporting structural correspondence at the level of tract geometry. Together, these findings underscore the translational value of the porcine model as a robust and neurobiologically relevant platform for investigating human brain function, structural organization, and related neurological disorders.

## Introduction

Cross-species comparative mapping of functional homologies provides a critical framework for understanding human cortical organization and function. Over the past several decades, studies in small animal models, particularly rodents, have substantially advanced our knowledge of brain structure, functional mechanisms, and the pathophysiology of neurological disorders [1–7]. Rodent models remain essential in preclinical neuroscience research [8–10]; however, significant translational gaps persist between preclinical findings and clinical application, largely due to fundamental differences in brain size, cytoarchitecture, and network organization between rodents and humans [1, 2, 11].

To address these limitations, parallel investigations have been conducted in nonhuman primates (NHPs), including marmosets, rhesus macaques, and chimpanzees, whose brains share closer anatomical and functional similarities with humans [12–14]. Comparative studies between humans and NHPs have demonstrated substantial functional correspondence and evolutionary conservation across multiple large-scale networks [15–18]. Nevertheless, ethical constraints, high costs, and practical limitations restrict the scope and scalability of experimental research in these species, thereby limiting further progress in elucidating human brain function [19, 20].

The porcine model has emerged as a compelling alternative for investigating human brain function and neurological disorders [23–26]. Compared with rodents, pigs possess a gyrencephalic cortex with cortical folding patterns more similar to those of humans, and their brain size more closely approximates the human brain [21–23, 27]. These anatomical features enhance their translational relevance (***Fig.1***). Furthermore, from both structural and functional connectomic perspectives, the pig provides a valuable model for investigating brain disorders compared to rodents and NHPs, particularly traumatic brain injury (TBI). Previous studies have demonstrated that porcine models yield clinically relevant insights into the structural and functional connectomic mechanisms underlying TBI, while also serving as a valuable platform for evaluating potential therapeutic interventions [28–30]. Beyond this, comparative neuroimaging studies have highlighted strong anatomical correspondence between human and porcine brains, including well-aligned white matter architecture [26, 31]. Therefore, porcine models have increasingly been utilized in translational neuroscience research, including studies of Huntington’s disease [32], Alzheimer’s disease [33], amyotrophic lateral sclerosis [34], Parkinson’s disease [35], and TBI [28–30, 36, 37].

**Figure 1:**
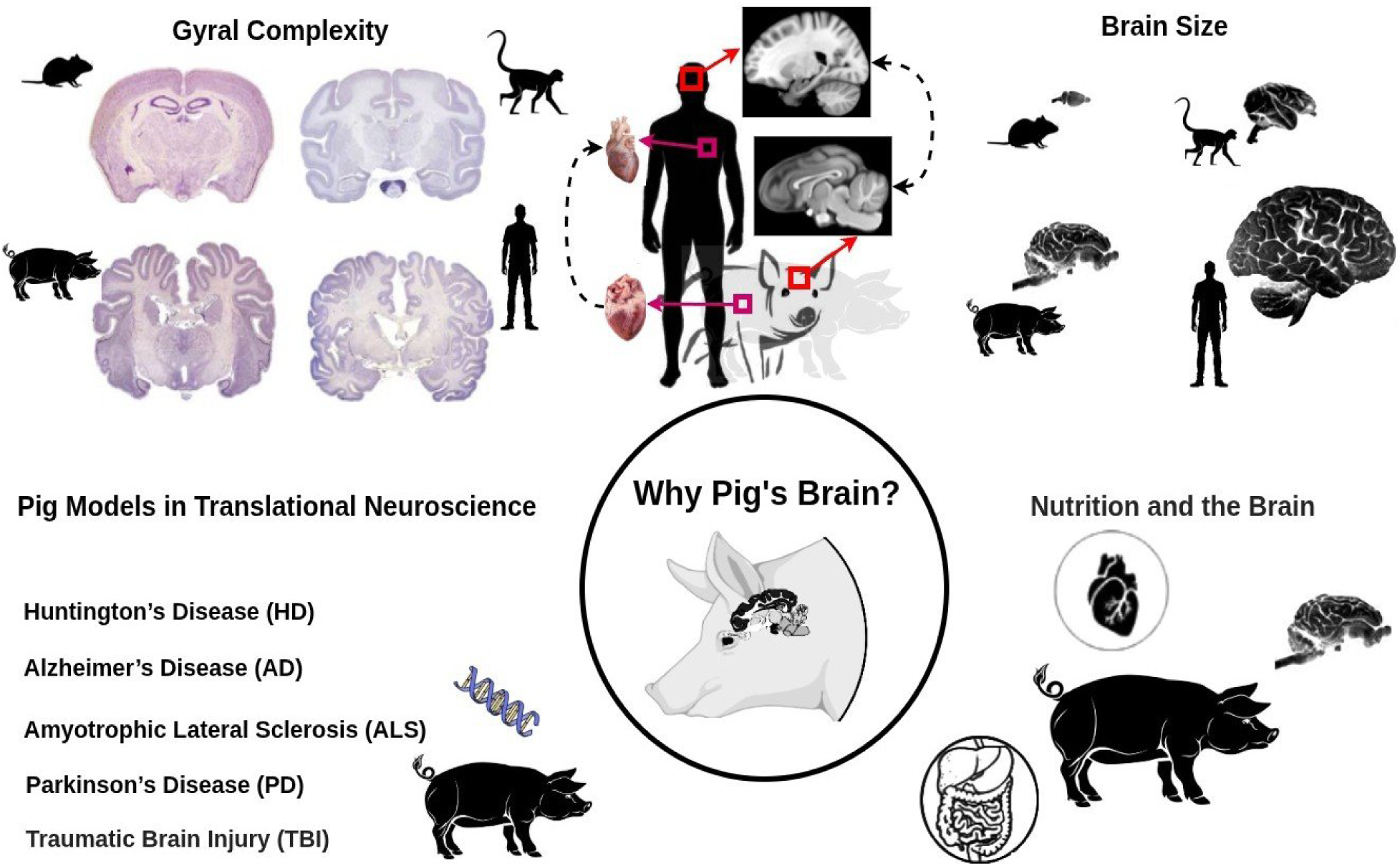
Why pig’s brain? Pigs exhibit greater gyral and structural similarity to humans compared with rodents and rhesus monkeys, including a more comparable brain size at a similar scale [21–23]. Porcine models are increasingly used in translational neuroscience, as well as in studies of nutrition, systemic health, and organ function beyond the brain. Therefore, investigating the neuroanatomy and functional organization of the porcine brain is essential for understanding human brain function and advancing translational research.

Beyond neuroscience, pigs play an important role in biomedical research more broadly [38–41]. Because pigs share metabolic and nutritional characteristics that closely resemble those of humans, they serve as a highly relevant animal model for investigating how dietary factors affect brain health and interact with systemic conditions such as cardiovascular and metabolic disorders [42–46]. Moreover, pigs have been extensively investigated in xenotransplantation research, particularly in the context of cardiac replacement therapies [47–49].

Altogether, comparative investigation of the porcine and human brain provides a valuable framework for understanding large-scale brain organization and the pathophysiology of neurological disorders.

In this study, we collected resting-state functional Magnetic Resonance Imaging (rs-fMRI) and diffusion Magnetic Resonance Imaging (MRI) data from 42 pigs and, for cross-species comparison, analyzed 100 unrelated human rs-fMRI datasets from the Brain Genomics Superstruct Project (GSP) [50] along with diffusion MRI data from the Human Connectome Project (HCP) [51]. The GSP human rs-fMRI dataset was selected because its repetition time (TR=3s) matches that of the pig rs-fMRI scans, facilitating cross-species functional comparisons. To identify and align large-scale intrinsic component networks (ICNs) across species, we applied group independent component analysis (ICA) separately to the pig and human rs-fMRI data. For structural comparisons, we performed fiber tractography, examined color-encoded fractional anisotropy (FA) maps, and analyzed structural connectivity to assess the white matter architecture in pigs and humans. Through this integrated functional and structural approach, we aimed to map brain networks across species, providing deeper insight into cross-species brain organization and enhancing our understanding of human brain function.

## Results

### Canonical Large-Scale Resting-State Functional Brain Networks in the Pig Brain

The rs-fMRI data were collected from pigs using a 3T MRI scanner with a repetition time of 3 seconds (***Fig.2A***). To identify functional networks, data from all subjects were combined over time and analyzed using group-level ICA, producing spatial maps of networks along with their associated temporal activity patterns (***Fig.2B***). Analysis of pigs’ rs-fMRI data identified twelve components from an initial set of seventy, which were selected based on spatiotemporal characteristics and visual inspection and grouped into seven large-scale functional networks (***Fig.2C***). These networks encompassed a broad range of cortical regions, including sensory, motor, visual, and associative areas, with each network exhibiting distinct spatial organization and minimal overlap with other networks.

**Figure 2:**
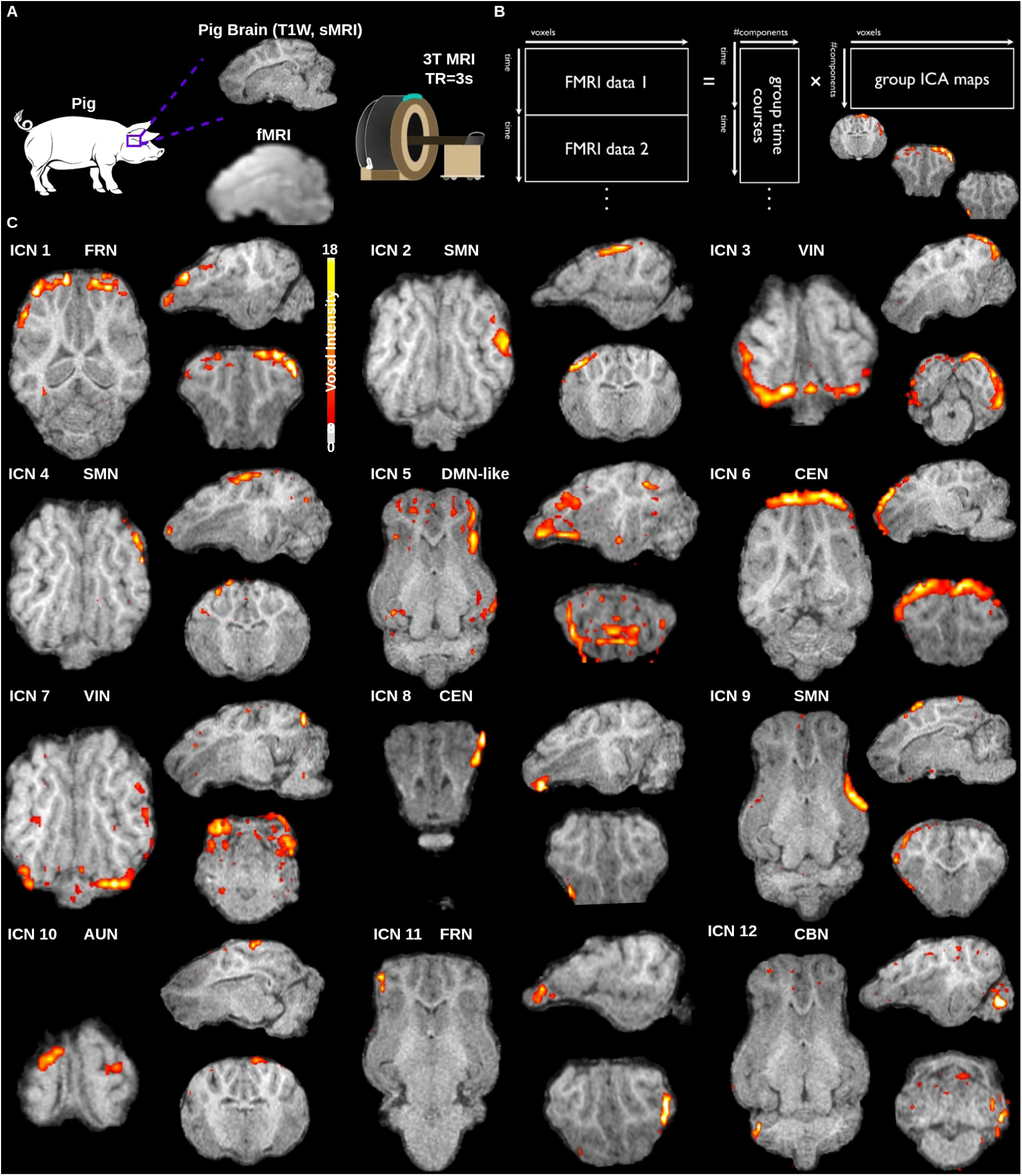
Whole-brain functional networks in the pig brain. **A.** Pig structural and rs-fMRI data were acquired using a 3T MRI scanner (rs-fMRI, TR=3s). **B.** Group ICA was performed on temporally concatenated data across pig subjects to derive spatial network maps and their corresponding temporal profiles. **C.** Identified components are displayed on sagittal, coronal, and axial slices of a high-resolution structural image of the pig brain. The networks include the frontal network (FRN), sensorimotor network (SMN), visual network (VIN), default mode network-like (DMN-like), cerebellar network (CBN), auditory network (AUN) and central executive network (CEN). Colors represent voxel-wise z-scores, thresholded at *p <* 0.05, reflecting the posterior probability that a voxel belongs to the active distribution rather than noise.

The frontal network (FRN; ICN1, ICN11) was confined to prefrontal cortical regions. The sensorimotor network (SMN; ICN2, ICN4, ICN9) spanned primary motor and somatosensory cortices. The visual network (VIN; ICN3, ICN7) was localized to occipital visual areas. The default mode network-like (DMN-like; ICN5) encompassed medial prefrontal, posterior cingulate, and lateral parietal cortices. The cerebellar network (CBN; ICN12) was restricted to cerebellar structures. The auditory network (AUN; ICN10) involved bilateral auditory cortices. The central executive network (CEN; ICN6, ICN8) included dorsolateral prefrontal and parietal cortical regions. Each network displayed consistent spatial patterns, and overlap between networks was minimal, with clear anatomical boundaries.

The pig brain exhibits a well-organized functional architecture, with discrete networks spanning sensorimotor, visual, and higher-order cortical regions, providing clear evidence that canonical large-scale resting-state networks are present. These networks show spatial patterns that closely correspond to those observed in humans. Given its gyrencephalic structure, the pig brain more closely resembles the human brain in anatomy, growth, and development than commonly used small laboratory animals such as rodents. As a translational large-animal model, pigs demonstrate resting-state networks that make them suitable for comparative and functional neuroscience studies.

### Canonical Large-Scale Resting-State Functional Brain Networks in the Human Brain

For comparison with the resting-state functional networks observed in the pig brain, we used a sample of human brain data from the GSP [50]. This dataset was chosen because it provides high-resolution, high-quality structural and functional MRI data (***Fig.3A***). In addition, the human fMRI data were acquired with a repetition time of 3 seconds, matching the acquisition parameters used for the pig rs-fMRI scans. The same processing pipeline was applied to the human data (***Fig.3B***), starting with seventy components, followed by visual inspection and selection. This procedure resulted in the identification of twelve resting-state ICNs in the human brain, which closely corresponded to large-scale networks observed in the pig brain, showing a strong spatial alignment between species.

**Figure 3:**
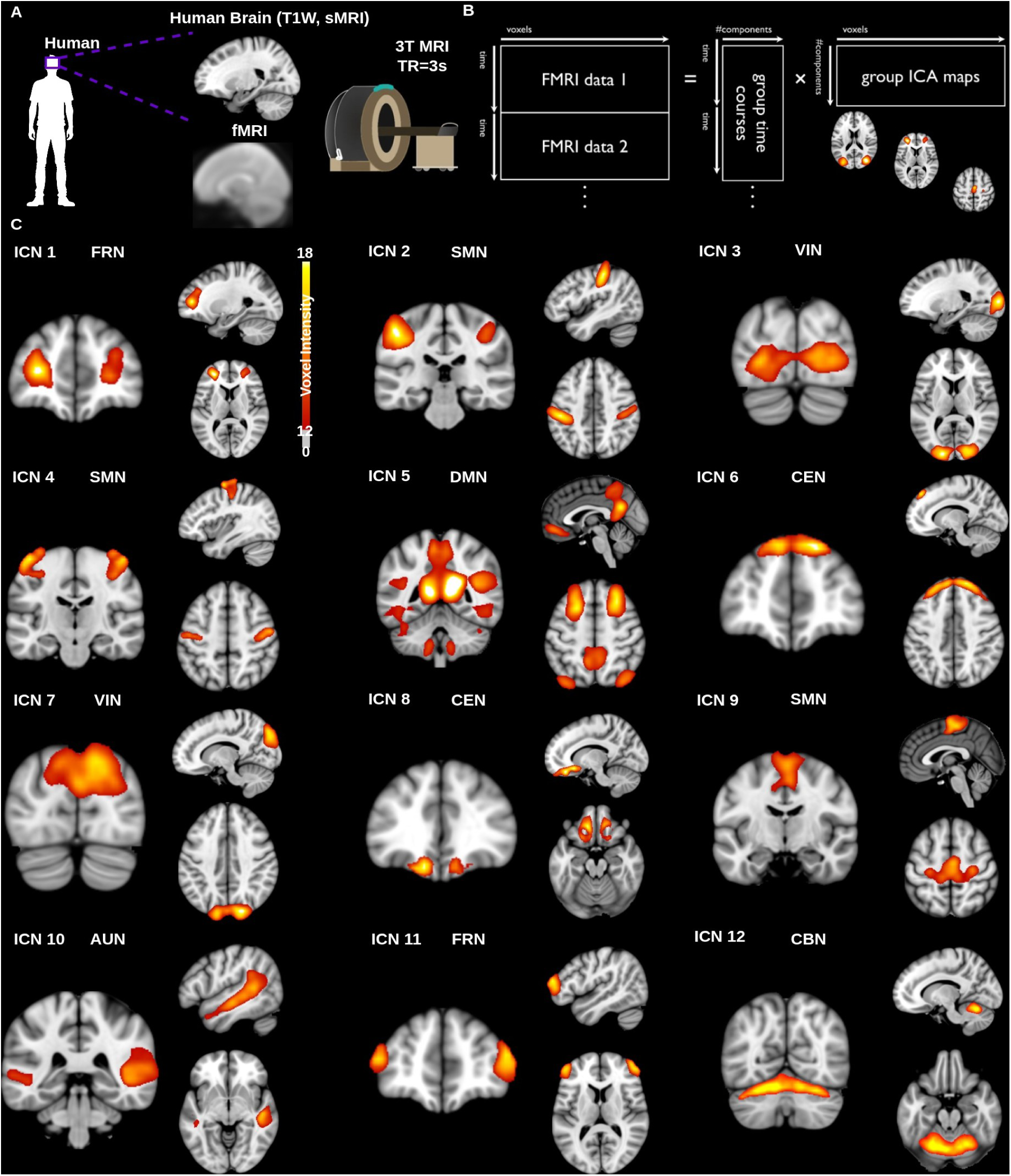
Whole-brain functional networks in the human brain. **A.** Human structural and rs-fMRI data were acquired using a 3T MRI scanner (rs-fMRI, TR=3s). **B.** Group independent component analysis was performed on temporally concatenated data across human subjects to derive spatial network maps and their corresponding temporal profiles. **C.** Identified components are displayed on sagittal, coronal, and axial slices of a high-resolution structural template image of the human brain. The networks include the frontal network (FRN), sensorimotor network (SMN), visual network (VIN), default mode network (DMN), cerebellar network (CBN), auditory network (AUN), and central executive network (CEN), which are consistent with the corresponding networks identified in the pig brain. Components were selected based on spatiotemporal characteristics and visual inspection.

**Figure 4:**
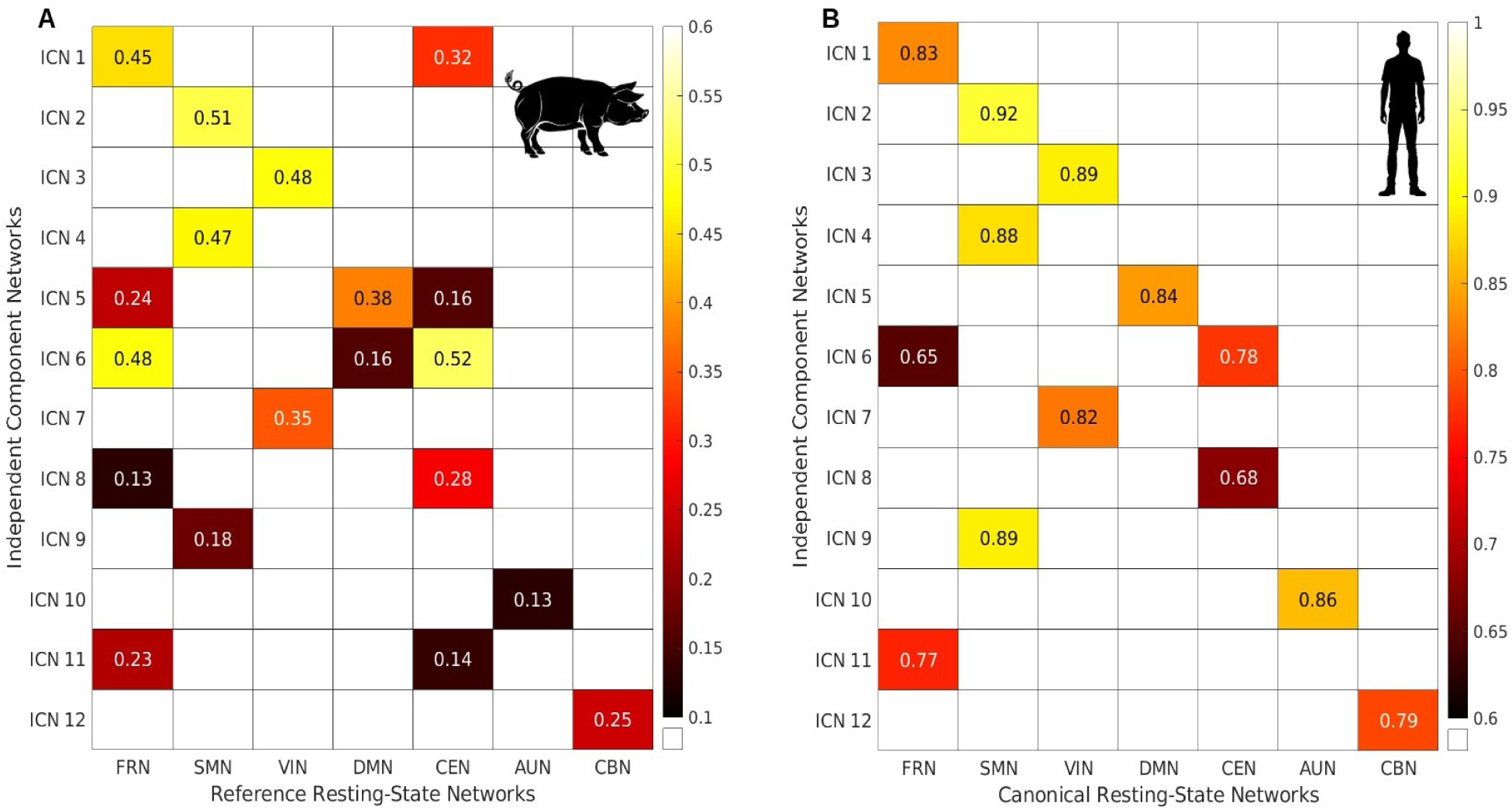
Spatial correlation between ICNs and reference (canonical) resting-state networks. **A.** Spatial correlation between pig ICNs and reference resting-state networks. **B.** Spatial correlation between human ICNs and canonical human resting-state networks.

**Figure 5:**
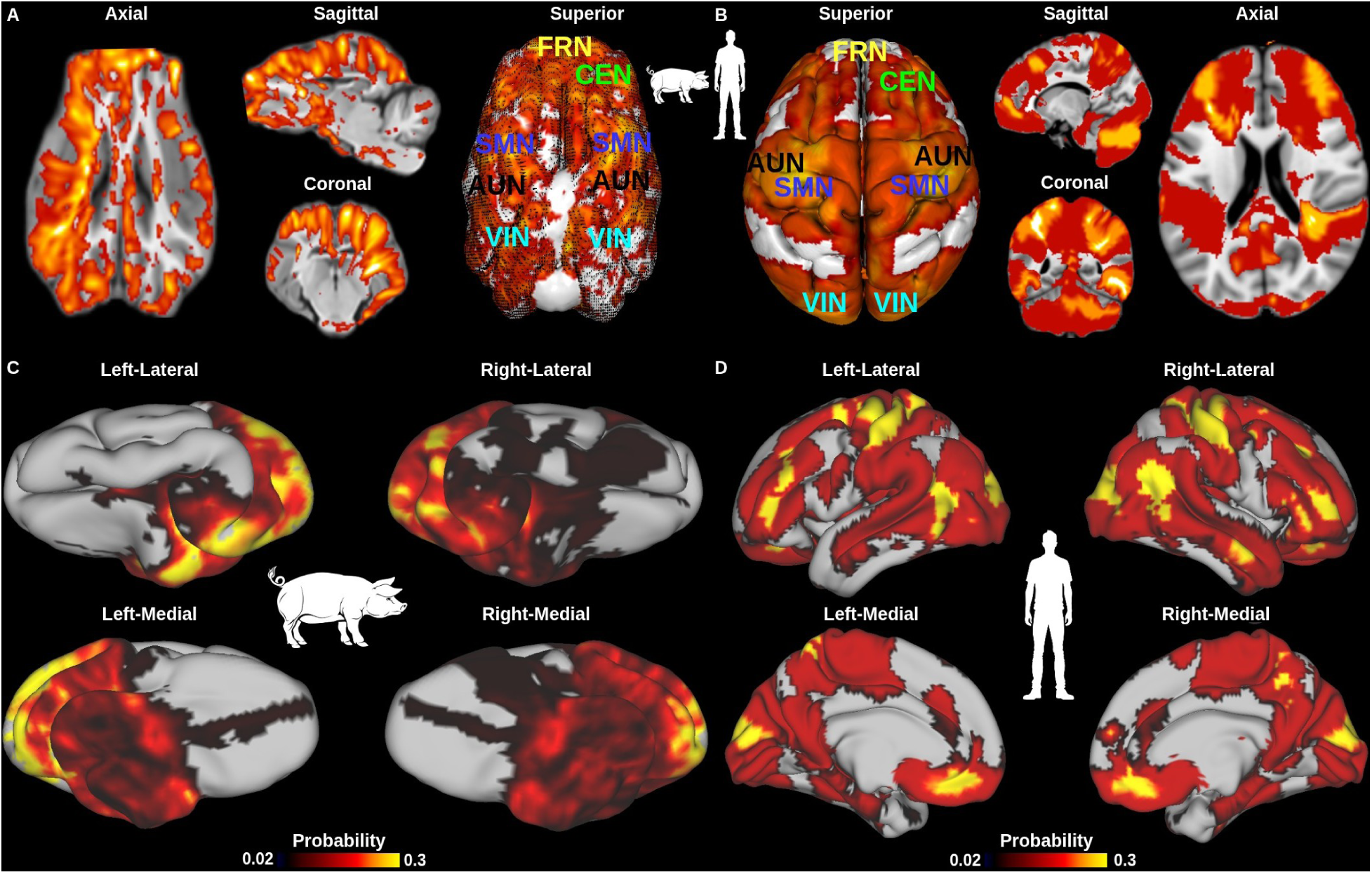
Probability maps of network engagement in pig and human brains. **A.** Voxel-wise volume probability map in pigs illustrating the frequency with which each voxel contributed to the identified functional components. Voxels with the highest recurrence were predominantly localized within the sensorimotor, prefrontal, visual, and cerebellar regions. Higher voxel-wise recurrence (probability) values are shown in yellow, whereas lower recurrence values are shown in black. **B.** Voxel-wise volume probability map in humans illustrating the frequency of voxel involvement across functional components. Highly recurrent voxels were predominantly observed in the prefrontal, sensorimotor, visual, and cerebellar regions, which closely align with those identified in pigs. **C.** Voxel-wise probability map displayed on inflated cortical surface renderings of the Porcine Neurological Imaging Space atlas [31]. **D.** Voxel-wise probability map visualized on inflated brain surface renderings of the Human Connectome Project Multi-modal atlas [52].

Twelve resting-state ICNs were identified and grouped into seven large-scale networks (***Fig.3C***). These included the frontal network (FRN; ICN1, ICN11), the sensorimotor network (SMN; ICN2, ICN4, ICN9), the visual network (VIN; ICN3, ICN7), the default mode network (DMN; ICN5), the cerebellar network (CBN; ICN12), the auditory network (AUN; ICN10), and the central executive network (CEN; ICN6, ICN8).

### Homologous Resting-State Networks in Pig and Human Brains

To investigate cross-species correspondence, rs-fMRI data from humans were analyzed alongside the pig data. Across both species, twelve independent components were identified and grouped into seven large-scale resting-state networks: the frontal network, sensorimotor network, visual network, default mode network, cerebellar network, auditory network, and central executive network. In both pigs and humans, these networks displayed distinct spatial patterns with minimal overlap, and their overall organization was consistent across subjects within each species. Comparison of the spatial maps revealed strong correspondence between pig ICNs and canonical human resting-state networks, indicating substantial conservation of large-scale functional network organization across species (***Figs.4A,B***). These results further support the use of pigs as a translational large-animal model for studying human brain function, providing a valuable platform for investigating human cognitive processes and neurological disorders.

To further quantify network organization, we examined the spatial distributions of temporal signal-to-noise ratio (tSNR) maps (***SI, Supplementary Figure S1***) and frequently recruited voxels in both pigs and humans, enabling a comparison of regional signal quality and network involvement across species. In the pig brain, these voxels were most prominent in prefrontal, sensorimotor, auditory, and visual cortices, as well as in the cerebellum (***Figs.5A,C***). Comparative analysis with the human brain revealed similar patterns of voxel recruitment across corresponding cortical regions, demonstrating a conserved topographical organization of resting-state networks between the two species (***Figs.5B,D***). Importantly, this correspondence was evident across multiple networks, including the frontal, sensorimotor, visual, and default mode networks, showing that pigs and humans share not only the same large-scale network architecture but also similar patterns of voxel recruitment within these networks. These findings underscore the conservation of functional topography and indicate that the spatial distribution of network activity is largely preserved across gyrencephalic brains.

### Temporal Correspondence of Functional Networks Between Pigs and Humans

To further quantify the temporal correspondence of functional networks in pigs and humans, we extracted the time courses from each identified network in both species (***Fig.6A***). Pearson correlation analyses were performed to quantify temporal similarity within species across networks and between species for corresponding networks.

**Figure 6:**
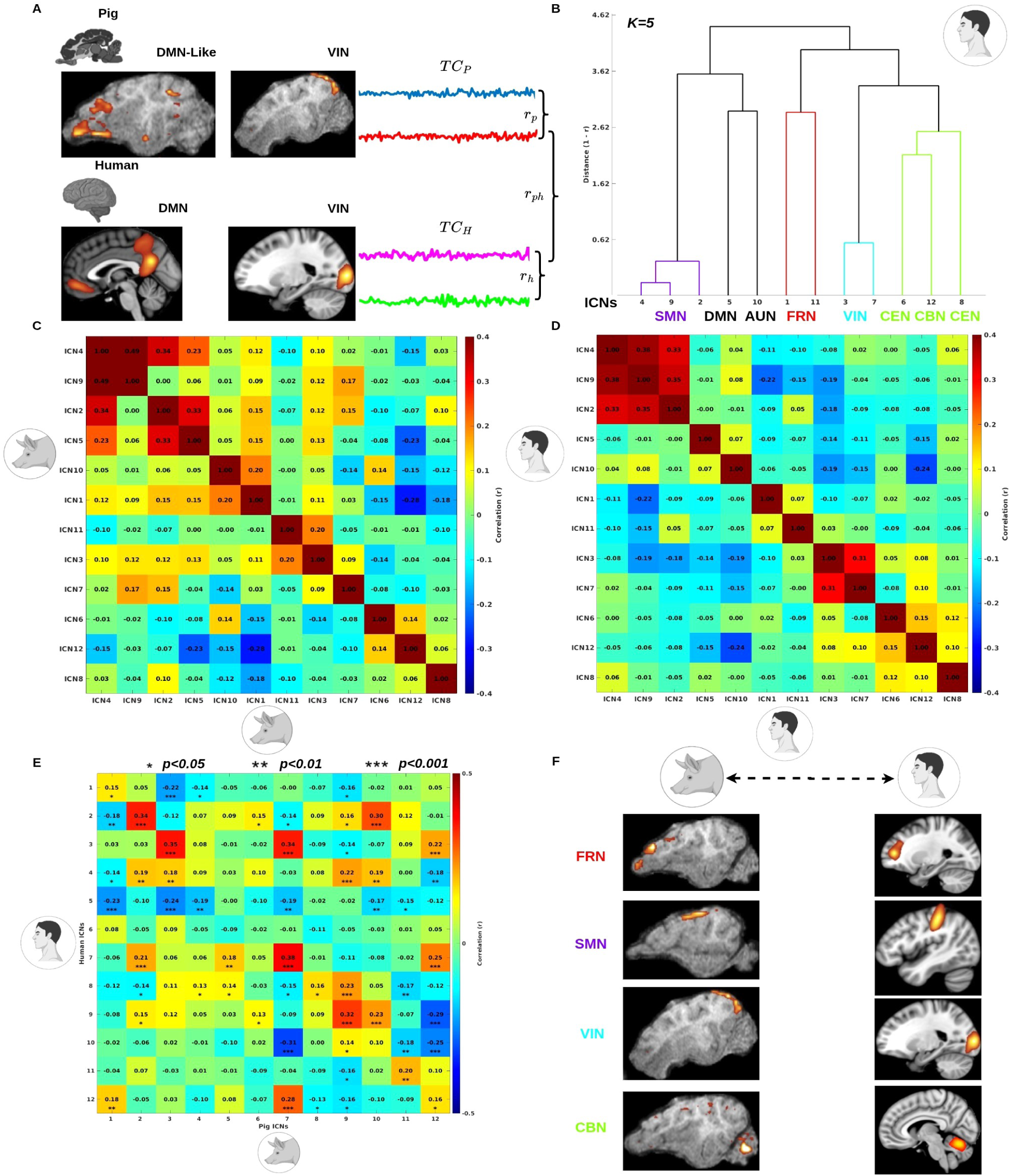
Comparison of within- and cross-species functional brain networks. **A.** To assess within- and cross-species functional similarity, the aligned networks and their corresponding time courses were extracted for computation. **B.** Hierarchical clustering dendrogram of the twelve functional human components using squared Euclidean distance and Ward’s linkage. Identified clusters include sensorimotor (purple), default mode and auditory (black), frontal (red), visual (cyan), central executive networks and cerebellar (green). **C.** Within-species correlation matrix for pigs showing pairwise temporal correlations (Pearson’s r) between component time courses. Components are ordered according to the hierarchical clustering tree. **D.** Within-species correlation matrix for humans showing pairwise temporal correlations between component time courses. Components are ordered according to the human hierarchical clustering tree. **E.** Matrix showing temporal correlations between twelve pig and twelve human functional components. Warm colors indicate positive correlations, while cool colors indicate anti correlations. Asterisks denote correlations that reached statistical significance after False Discovery Rate (FDR) correction based on a permutation test (\**p <* 0.05, \*\**p <* 0.01, \*\**p <* 0.001). **F.** Representation of networks exhibiting high and significant correlations between pigs and humans.

To further characterize the organization of network relationships, hierarchical clustering was applied to the human correlation matrix to identify groups of temporally similar networks (***Fig.6B***). Based on this clustering structure, the correlation matrices were reordered accordingly and the same ordering was applied to the pig data (***Figs.6C,D***). The reordered matrices revealed comparable clustering patterns in both species, with functionally related networks exhibiting stronger within-cluster correlations. In particular, the sensorimotor, visual, frontal, and central executive networks formed coherent clusters characterized by higher internal temporal similarity relative to correlations with other networks. This consistent clustering structure across pigs and humans indicates a similar temporal organization of large-scale functional networks.

Cross-species analysis also revealed significant positive correlations between homologous networks, demonstrating similar temporal fluctuation patterns in pigs and humans (***Fig.6E***). Corresponding networks showed higher interspecies correlations compared to non-matching networks, indicating specificity in temporal alignment. This effect was particularly evident in the frontal, sensorimotor, visual, and cerebellar networks, which demonstrated stronger cross-species temporal similarity relative to other network pairings (***Fig.6F***). These findings indicate that homologous networks not only share similar spatial organization but also exhibit aligned temporal dynamics across pigs and humans.

Together with the spatial correspondence observed earlier, these findings show that both the large-scale topography and the intrinsic temporal dynamics of resting-state networks are preserved across the two species.

### Comprehensive White Matter Tractography of the Pig Brain

To investigate white matter organization in the pig brain, diffusion-based fiber tractography was performed to reconstruct structural connections across the whole brain. The resulting tractography provided a comprehensive representation of large-scale white matter architecture, revealing extensive intrahemispheric and interhemispheric fiber pathways throughout cortical and subcortical regions (***Fig.7A***).

**Figure 7:**
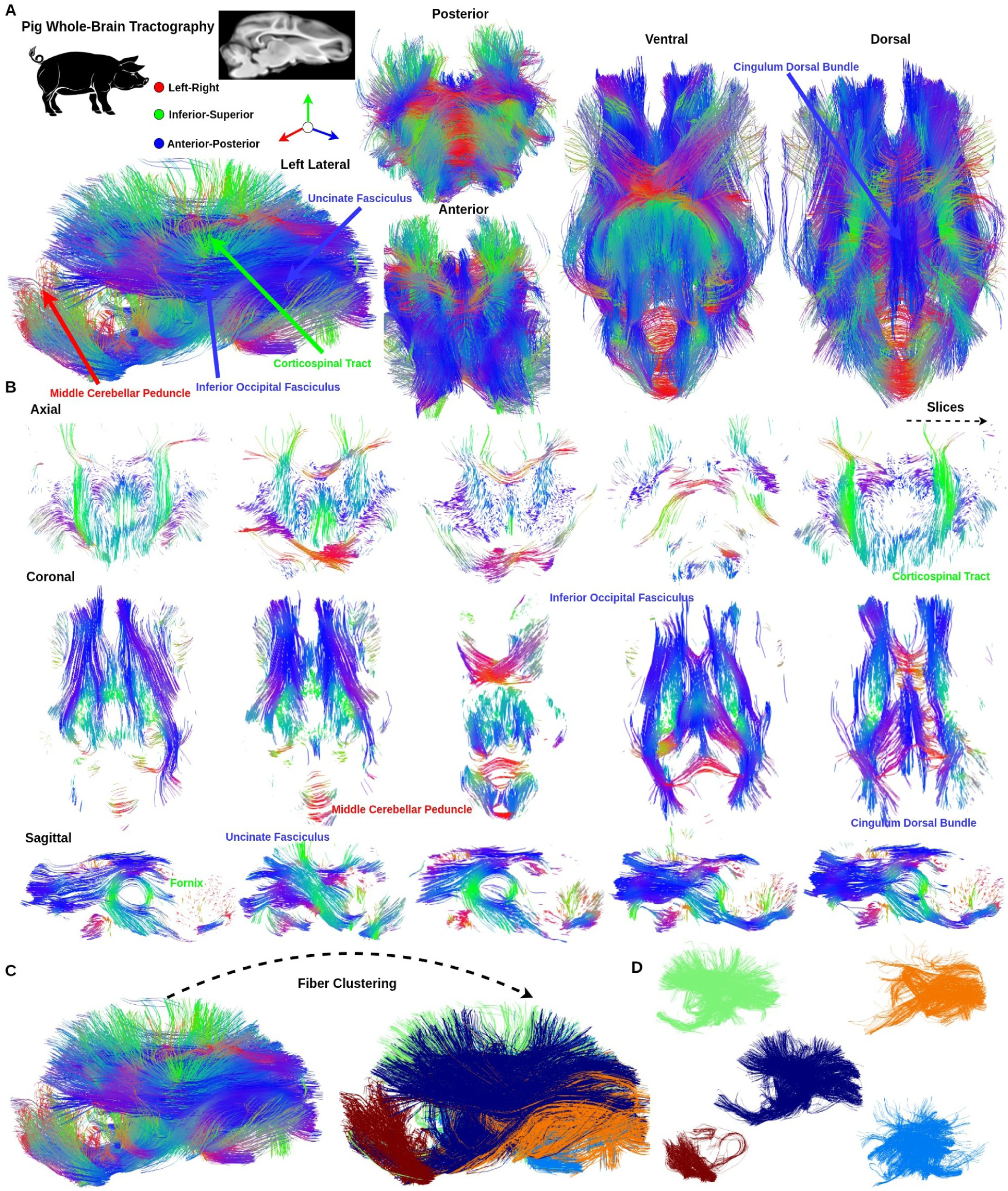
Pig whole-brain tractography. **A.** Whole-brain tractography of the pig was generated from diffusion MRI. The fiber tracts are illustrated from multiple views (left lateral, posterior, anterior, ventral, and dorsal) with fibers color-encoded according to orientation. **B.** The fiber tracts from the slice series are visualized in axial, coronal, and sagittal views, with fibers color-encoded according to orientation to facilitate tracking of fiber trajectories. **C.** Fiber clustering was applied to group fibers into distinct families, resulting in the identification of five clusters. **D.** The five fiber clusters are presented using distinct colors.

To clearly visualize fiber orientation in the whole pig brain, fiber tractography was examined in axial, coronal, and sagittal views (***Fig.7B***). These multi-planar representations allowed detailed observation of major projection, association, and commissural fiber systems. Notable tracts, including the Inferior Occipital Fasciculus, Corticospinal Tract, Uncinate Fasciculus, Fornix, Cingulum Dorsal Bundle, and Middle Cerebellar Peduncle, were distinctly delineated, illustrating the overall structural connectivity and organization of the pig brain (***SI, Supplementary Figures S2, S3***).

To further investigate the fine structure of the pig brain, fiber clustering was applied to the whole-brain tractography data. This analysis identified five major fiber clusters, representing the primary structural organization of pig brain white matter (***Fig.7C***). Each cluster corresponded to a distinct set of projection, association, or commissural pathways (***Fig.7D***). Notable examples included the Middle Cerebellar Peduncle (deep red), Corticospinal Tract (green), Anterior Thalamic Radiation (light blue), Uncinate Fasciculus (orange), and Inferior Occipital Fasciculus (deep blue). Together, these clusters captured the major topography and connectivity patterns of the pig brain’s primary fiber systems, providing a comprehensive overview of structural organization.

To quantify the structural organization of white matter in the pig brain, structural connectivity (SC) matrices were generated from whole-brain tractography. Representative SC matrices from three pigs (Pig 1, Pig 12, and Pig 38) were selected as examples, as they exhibited highly similar sparse connectivity patterns (***Fig.8A***). In addition, a group-average SC matrix was computed across all 42 pigs (***Fig.8B***). Pairwise comparisons revealed a high degree of similarity between individual SC matrices, with all inter-subject correlations reaching statistical significance (FDR-corrected *p <* 0.001) (***Fig.8C***). Specifically, the correlation coefficients were *r* = 0.941 for Pig 1 vs. Pig 12, *r* = 0.925 for Pig 1 vs. Pig 38, and *r* = 0.921 for Pig 12 vs. Pig 38. Furthermore, each individual SC matrix showed strong agreement with the group-average SC matrix (***Fig.8D***). For example, correlations with the average SC matrix were *r* = 0.965 for Pig 1, *r* = 0.964 for Pig 12, and *r* = 0.964 for Pig 38 (all FDR-corrected *p <* 0.001).

**Figure 8:**
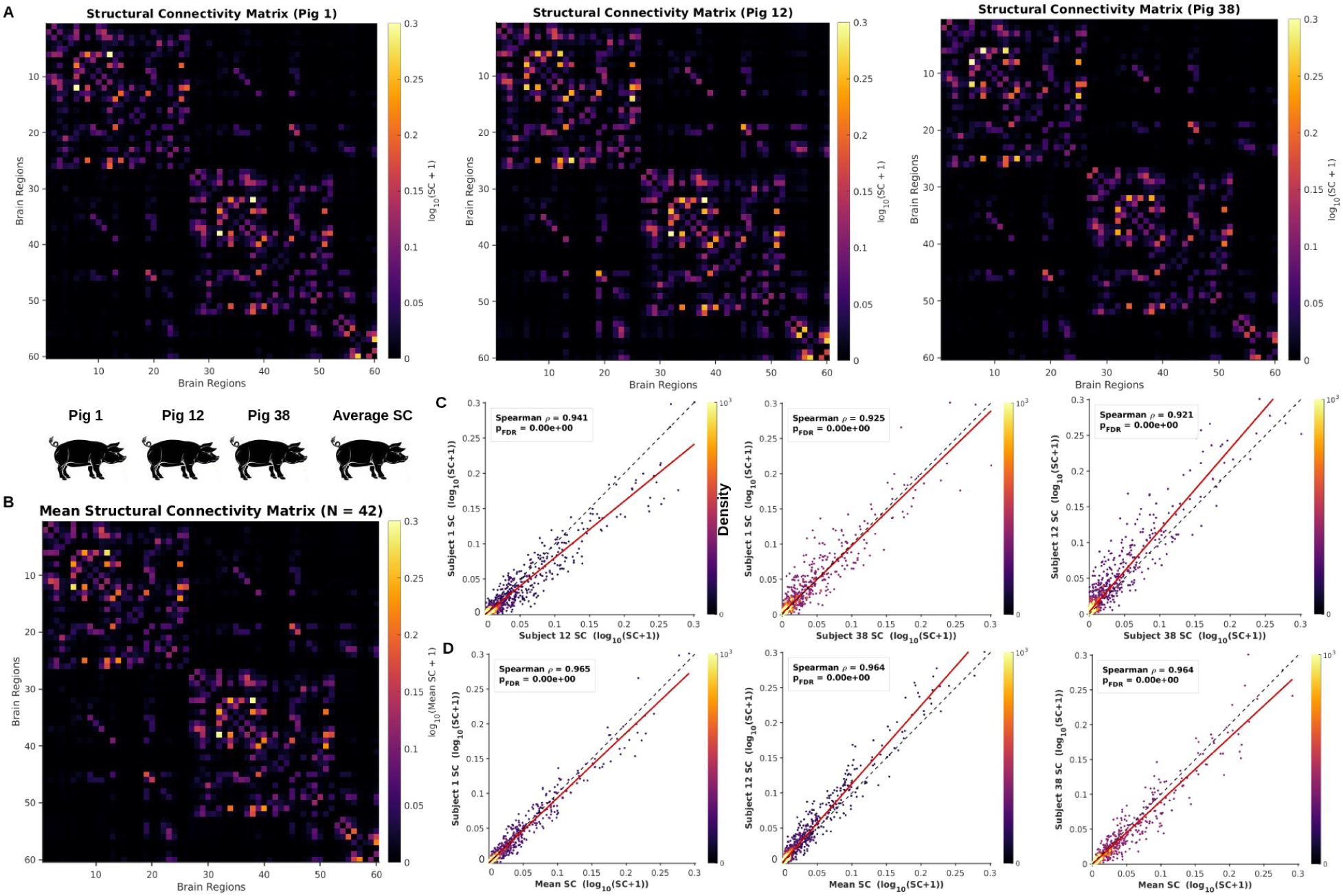
Structural connectivity of the pig brain. **A.** Representative whole-brain structural connectivity matrices from Pig 1, Pig 12, and Pig 38. Matrix entries represent structural connection strengths between brain regions and are displayed as log_10_(SC + 1) to enhance visualization. **B.** Average structural connectivity matrix across all pigs (*n* = 42). **C.** Inter-subject similarity of structural connectivity matrices among representative pigs, along with the corresponding correlation coefficients (*r*) and associated FDR-corrected *p*-values. **D.** Correlation between each individual pig’s structural connectivity matrix and the group-average structural connectivity matrix.

Taken together, these results demonstrate a highly reproducible structural connectivity architecture across pigs, indicating that the tractography-derived connectomes capture consistent and robust brain-wide SC patterns at both the individual and population levels.

### Conserved Whole-Brain White Matter Architecture Across Pigs and Humans

To assess comparative structural architecture across pigs and humans, dense and sparse whole-brain fiber tractography was constructed for both species, capturing white matter organization at multiple levels of detail. The dense tractography captured the complete white matter architecture (***Figs.9A,D***), while the sparse representation highlighted the major fiber bundles (***Figs.9B,C***).

In addition, color-encoded FA maps were generated for both pigs and humans to visualize white matter microstructure and fiber orientation (***Fig.9E***). In both species, major white matter tracts were clearly delineated, with consistent orientation patterns observable across corresponding regions. These maps revealed that projection, association, and commissural fibers exhibited similar directional organization in pigs and humans, supporting the presence of homologous white matter architecture. The color-encoded FA representation allowed direct visual comparison of tract trajectories and overall structural layout, highlighting both conserved features and species-specific differences in fiber orientations and tract prominence.

**Figure 9:**
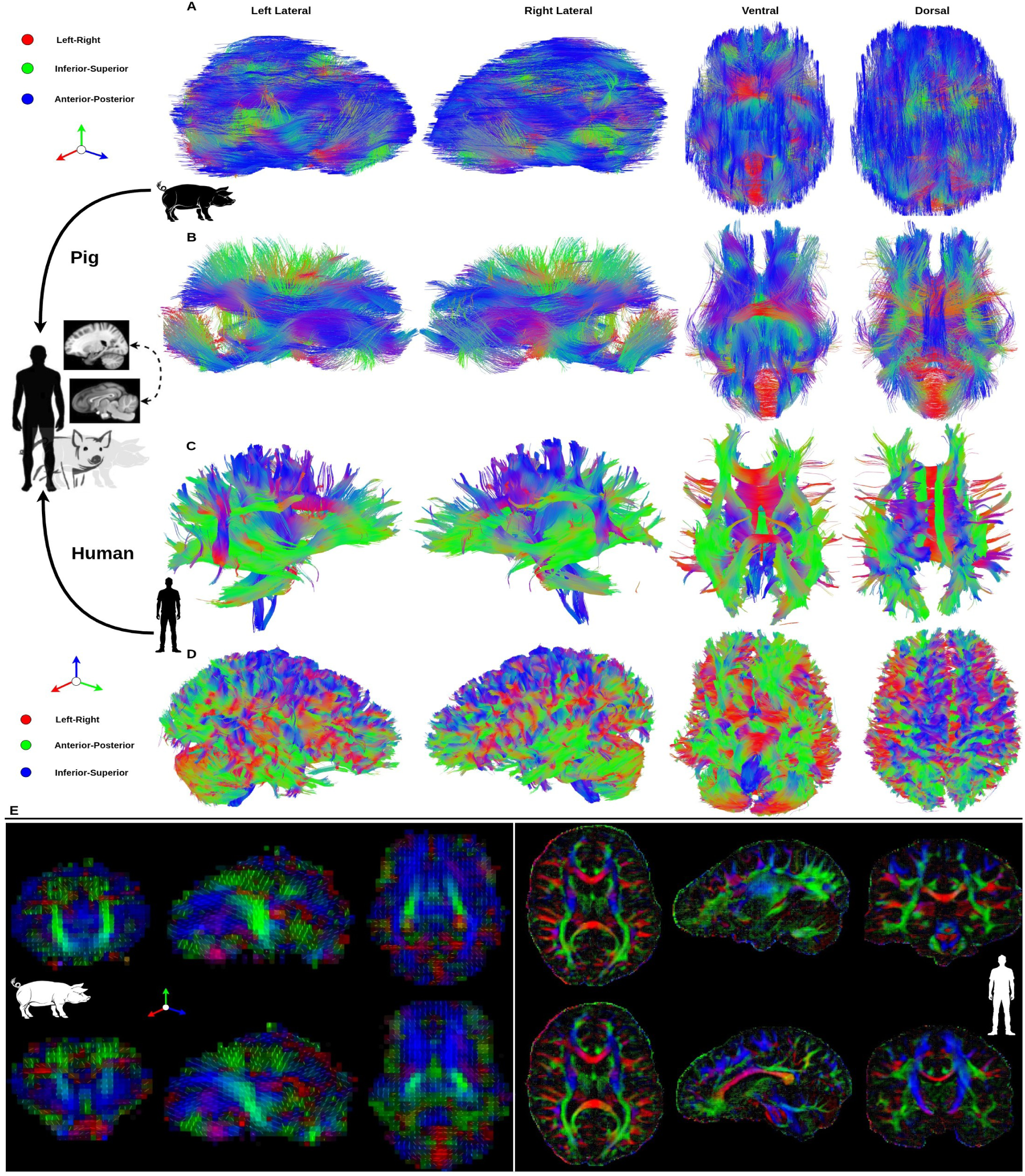
Cross-species comparison of whole-brain tractography in pigs and humans. **A.** Dense whole-brain tractography in the pig was reconstructed from diffusion MRI data. The resulting fiber tracts are visualized from multiple perspectives, including left lateral, right lateral, ventral, and dorsal views, with fibers color-coded according to orientation. **B.** Sparse whole-brain tractography in the pig was reconstructed from diffusion MRI data to better track fiber orientations. The resulting fiber tracts are visualized from multiple perspectives, including left lateral, right lateral, ventral, and dorsal views, with fibers color-coded according to orientation. **C.** Sparse whole-brain tractography in humans was reconstructed from diffusion MRI data. **D.** Dense whole-brain tractography in humans was reconstructed from diffusion MRI data. **E.** Color-encoded FA maps were generated for both pigs and humans to facilitate comparison of white matter

**Figure 10:**
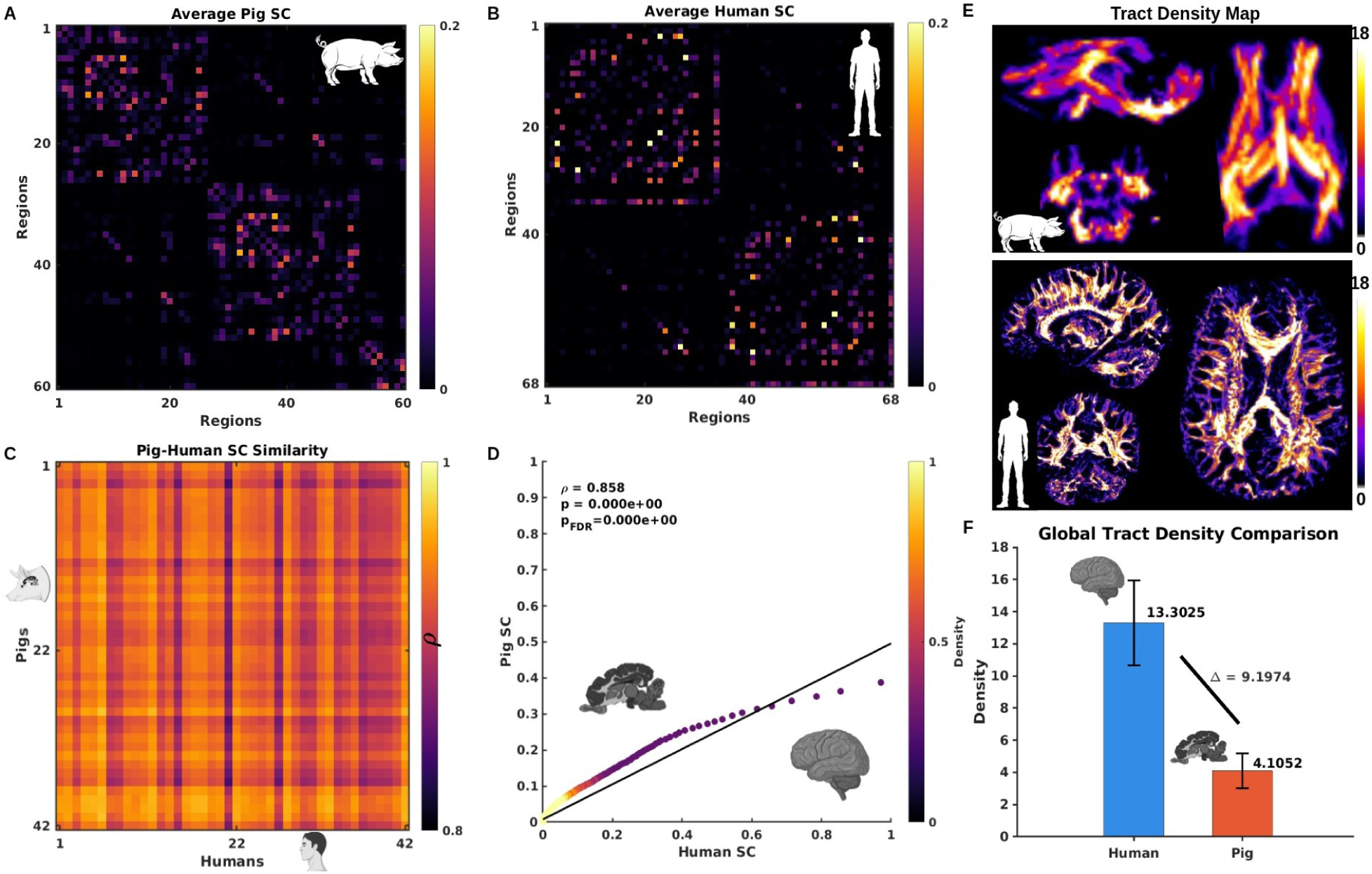
Comparison of structural connectivity between pigs and humans. **A.** Average structural connectivity matrix across all pigs (*n* = 42). **B.** Average structural connectivity matrix across all humans (*n* = 42). **C.** Pig-human similarity of structural connectivity matrices across all subjects. **D.** Correlation between each individual pig’s structural connectivity matrix and each human structural connectivity matrix, with corresponding correlation coefficients (*r*) and FDR-corrected *p*-values. **E.** Tract density maps for pigs and humans. **F.** Comparative tract density maps of pigs and humans. The normalized mean tract density value for each species is presented, together with the difference in mean tract density (Δ) between pigs and humans.

To further evaluate the conservation of white matter architecture across species, we compared the SC patterns between pig and human brains. Group-average SC matrices were first estimated for both pigs (***Fig.10A***) and humans (***Fig.10B***), and cross-species similarity analyses were subsequently performed (***Fig.10C***). The results revealed a high degree of similarity between pig and human SC organization, with an average correlation coefficient of *r* = 0.858 (FDR-corrected *p <* 0.001) (***Fig.10D***). In addition, we also comparative analysis revealed that humans exhibited higher overall fiber density relative to pigs (***Figs.10E,F***), reflecting more extensive large-scale white matter organization. Despite this difference in density, the overall structural layout was highly conserved between species, with homologous projection, association, and commissural pathways clearly identifiable in both pigs and humans. These findings suggest that the large-scale organization of white matter connectivity is highly conserved between pigs and humans, supporting the translational value of the pig brain as a model for investigating human brain structural networks.

### Conserved Functional and Structural Brain Architecture in Pigs and Humans

Through neuroimaging, we performed rapid and comprehensive phenotyping of cortical organization in pigs and humans. Both species exhibited highly similar functional and structural brain organization. Resting-state fMRI analysis revealed homologous large-scale networks, such as the frontal, sensorimotor, and visual networks, with comparable spatial and temporal properties. White matter organization, assessed using fiber tractography and structural connectivity analysis, revealed highly similar patterns in both species. Homologous white matter tracts, including the corticospinal tract and uncinate fasciculus, were clearly delineated, and the overall structural connectivity architecture was highly consistent between pigs and humans. These findings indicate that the global white matter architecture is largely conserved across species, further underscoring their strong structural and functional similarity.

## Discussion

Pigs are emerging as valuable biomedical models, particularly in neuroscience, serving as large-animal models for the study of neurological diseases [53, 54]. Despite earlier efforts to investigate functional and structural homology between pig and human brains [26, 28, 31, 55], these relationships remain largely unexplored and poorly understood.

In this study, we examined the organization of pig and human brains, considering both functional and structural level. Our results demonstrated that pigs and humans share highly similar large-scale functional networks, including the frontal, sensorimotor, visual, and default mode networks, with comparable spatial and temporal patterns. Structural analyses using fiber tractography and structural connectivity further demonstrated that the overall white matter architecture is largely conserved between pigs and humans.

From a functional perspective, large-scale resting-state brain networks identified in the pig brain consistently align with canonical resting-state networks in humans. This correspondence provides a more comprehensive understanding of functional brain organization in pigs and highlights the presence of homologous network structures across species. The similarity in both spatial and temporal patterns of these networks suggests that key aspects of large-scale brain organization are conserved, reinforcing the translational relevance of pigs as a model for human neuroscience. Moreover, these findings support the use of pigs for investigating functional connectivity alterations in neurological disorders [24, 29, 36, 56, 57], as the homologous networks provide a framework to relate findings directly to human brain function.

However, our cross-species analysis of temporal correspondence focused on pairwise correlations, providing a limited view of brain network interactions. As previous studies have suggested [58–61], brain connectivity extends beyond conventional pairwise relationships between brain regions. Functional interactions can also emerge among groups of three or more regions simultaneously, including triadic, tetradic, and higher-order dependencies. These multivariate interactions, commonly referred to as high-order functional interactions, capture coordinated activity patterns that cannot be fully explained by pairwise correlations alone. As such, they are critical for characterizing the complexity of brain network organization and revealing integrative mechanisms underlying large-scale neural communication.

Therefore, future studies should examine higher-order functional connectivity to enable more comprehensive cross-species comparisons. Additionally, recognizing the dynamic nature of functional connectivity [62], rather than treating neural interactions as static, may yield deeper insight into information processing in the pig brain and further strengthen its translational relevance to human neuroscience.

Furthermore, it is important to note that pig brain imaging was performed under general anesthesia, whereas human rs-fMRI data were collected from awake participants. Anesthesia is known to alter neuronal activity and overall brain network dynamics, which can influence measures of functional connectivity and whole-brain organization [63, 64]. Consequently, differences in brain activity between anesthetized pigs and awake humans may introduce variability in cross-species analyses. Some discrepancies in network architecture or connectivity patterns could therefore reflect the effects of anesthesia rather than the inherent species-specific differences. This point should be considered when interpreting cross-species comparisons, particularly in studies examining functional brain network correspondences between pigs and humans.

From a structural perspective, diffusion-based tractography revealed that the overall organization of white matter pathways is highly conserved between pigs and humans. Major tracts, including the Corticospinal Tract, Uncinate Fasciculus, and Middle Cerebellar Peduncle, exhibited homologous trajectories in both species. While humans showed higher fiber density, the global layout and topography of projection, association, and commissural pathways were remarkably similar, supporting the notion of conserved structural architecture across gyrencephalic brains. In addition, we estimated structural connectivity matrices from tractography-derived fiber pathways in both pigs and humans to provide a detailed, quantitative assessment of white matter organization [65]. This analysis revealed a high degree of conservation in structural connectivity patterns across the two species, with an approximate similarity of 85%, which revealed a high degree of conservation in structural connectivity patterns between pigs and humans.

In future studies, integrating structural connectivity matrices with functional network analyses may help elucidate structure-function relationships, thereby providing a more comprehensive understanding of how anatomical connectivity constrains and supports coordinated brain activity [66].

One limitation of the current study is that pig fMRI data exhibited lower tSNR compared to human data. This reduced tSNR could arise from biological factors, such as differences in cortical thickness or vascular properties, as well as from technical factors related to scanning parameters or coil sensitivity. Lower tSNR may affect the sensitivity to detect subtle functional connectivity patterns and could contribute to underestimation of network strength. Future studies could improve tSNR by optimizing acquisition protocols, using higher-field scanners, or employing specialized coils designed for the pig brain to enhance the quality of functional imaging data [67, 68].

Another limitation of the study is that diffusion weighted imaging in pigs was acquired at a relatively lower spatial resolution (2 mm isotropic voxels) compared to the human scan (1.25 mm isotropic voxels). As a result, the resolution of the FA maps was lower in pigs, potentially limiting the fine-grained characterization of white matter microstructure. Employing higher spatial resolution in future pig studies could provide more detailed maps of white matter pathways and enable a more precise assessment of structural connectivity. Additionally, advanced diffusion models such as neurite orientation dispersion and density imaging (NODDI) could offer the opportunity to index microstructural properties and neuroplastic changes at the level of dendrites and synapses, further enhancing the translational value of pig models for studying human brain structure and function [69, 70].

In addition to data processing limitations, there are challenges associated with mapping pig brain data to the PNI50 surface atlas [31]. Volume-based probability maps in PNI50 space did not align as precisely to the cortical surface as in human studies, which may reduce the accuracy of spatial localization and cross-subject comparisons. Future studies could enhance surface mapping by refining the atlas, acquiring higher-resolution anatomical scans, or applying advanced registration techniques, allowing more accurate functional analyses in pigs.

In summary, this study represents an important first milestone in systematically comparing pig and human brain homology from both functional and structural perspectives. Future work could extend these analyses to investigate age-related trajectories of brain development. Such studies could examine how structural and functional connectivity in pigs corresponds to human brain maturation from early childhood through adolescence, providing valuable insights into conserved and species-specific neurodevelopmental patterns. These comparative developmental analyses could also inform translational models for pediatric neurological research and improve our understanding of the evolution of brain networks across mammals. In addition, based on our data and existing pig TBI models, we can leverage cross-species comparisons to investigate how TBI affects structural and functional connectivity during development. Such studies could provide translational insights into the mechanisms of pediatric brain injury and help guide the development of age-appropriate interventions and therapies.

## Materials and Methods

### Pigs

All procedures were approved by the Institutional Animal Care and Use Committee at the University of Georgia (Animal Use Protocol: A2023 07-021-Y3-A12) and adhered to the National Institutes of Health Guide for the Care and Use of Laboratory Animals, ensuring the ethical and humane treatment of all animals throughout the study.

Subjects in this study included 42 piglets (Sus scrofa domesticus, Yorkshire crossbreed), whose MRI scans were acquired at the Bio-Imaging Research Center, University of Georgia. The cohort consisted of 23 females and 19 males. Each animal was assigned a unique subject identifier, and sex information was recorded for all individuals.

To ensure safe and stress-free transportation, pigs will be given ondansetron (PO, 4mg tablet) and midazolam (IM, 0.4 mg/kg) immediately prior to being transported in mobile pig crates equipped with wheels and loaded into a temperature-controlled van. Upon arrival at the MRI facility, each animal will receive sedation consisting of xylazine (IM, 2mg/kg) and butorphanol (IM, 0.2mg/kg). Pigs will then be intubated and maintained with 1-3% isoflurane in oxygen. Throughout the MRI scans, continuous monitoring of temperature, respiration, and heart rate will be conducted. Once MRI is completed, the pig will be extubated, recovered, and then transported back to the pig facility.

### Humans

Human rs-fMRI datasets were obtained from the GSP [50], and diffusion weighted images were acquired from the HCP [51]. Both datasets are publicly available, open-access resources with ethical approval and consent protocols in place. For the present study, we used rs-fMRI data from 100 GSP subjects and diffusion weighted images from 42 HCP subjects to perform the following cross-species analyses. Full scanning parameters and preprocessing details for pig and human MRI data were provided in the ***Supplementary Materials and Methods***.

### Group ICA for Decomposition of Pig and Human Brain Activity

Group ICA [71] was performed to identify common functional networks in both pig and human datasets. This approach allows the decomposition of rs-fMRI data into spatially independent components (ICs), facilitating cross-species comparisons of functional brain organization.

For both species, participant-specific principal component analysis (PCA) was first performed to standardize the data and ensure that each participant contributed comparably to the shared subspace. This preprocessing step also improves computational efficiency and reduces noise by retaining the most informative variance components [72]. For each participant, the top 100 principal components (PCs) explaining the greatest amount of variance were retained for subsequent analyses. The retained PCs from all participants were then concatenated along the temporal dimension, and a second-level group PCA was conducted on the combined dataset to identify group-level variance patterns. While participant-specific PCA preserves individual variability, group-level PCA captures shared features across participants [73].

The top 80 group-level PCs explaining the maximum variance were subsequently used as inputs for group ICA. A model order of 70 components was selected based on previous studies and to enable the identification of large-scale functional brain networks. To evaluate the stability and reliability of the ICA decomposition, the Infomax algorithm implemented in ICASSO was used [74]. The ICA procedure was repeated 100 times with random initialization and bootstrapped fMRI time-series samples to generate multiple independent component solutions. Components that showed the highest similarity to the centroids of stable clusters were selected as the representative components for further analysis.

Component reliability was quantified using the ICASSO quality index (IQ), which measures the consistency of component estimation across repeated ICA runs [75]. Components were considered reliable large-scale brain networks if they met predefined criteria, including an IQ value greater than 0.80, substantial spatial overlap with gray matter, peak component weights localized within gray matter regions, and minimal spatial similarity to motion-related, ventricular, or other artifact components. Following network identification, the resulting independent components were classified based on their spatial activation patterns, and functional network labels were assigned according to established interpretations of their neurobiological roles. For both species, twelve independent component networks (ICNs) were identified and aligned across species. These components were then grouped into seven canonical functional networks.

### Spatial Correlation Analysis of Identified ICNs with Canonical Resting-State Networks in Both Species

To assign the identified ICNs in the pig brain to canonical resting-state networks (RSNs), we followed the same procedure described previously [26]. A reference pig RSN atlas was first constructed using the anatomical Saikali atlas [76], which was spatially normalized to the anatomical space of the template pig brain. To quantify the degree of homology between the group ICA-derived pig ICNs and established human RSNs, anatomically homologous pig brain regions corresponding to six major human RSNs [77] were manually identified and combined to construct six reference pig RSNs (***SI, Supplementary Łable S1***): the executive control network (CEN), cerebellar network (CBN), visual network (VIN), sensorimotor network (SMN), auditory network (AUN), frontal network (FRN), and default mode network (DMN). These manually constructed pig RSNs served as the reference atlas for evaluating the spatial correspondence of the group ICA-derived pig ICNs. Pearson’s spatial correlation coefficient was computed between each identified pig ICN and each reference pig RSN. For visualization, only correlation coefficients between 0.1 and 0.6 were displayed, while all other values were set to zero.

For the human data, the identified ICNs were assigned by computing their spatial correspondence with the NeuroMark 2.2 canonical RSN template [78], a multiscale functional network atlas derived from a large cohort of individuals. Pearson’s spatial correlation coefficient was calculated between each ICA-derived human ICN and each canonical RSN in the NeuroMark 2.2 template. For visualization, only correlation coefficients greater than 0.6 were retained, whereas all remaining values were set to zero.

### Cross-Species Comparison of Spatial and Temporal Correspondence

We computed within- and between-species correlation matrices of component time courses. To evaluate the statistical significance of cross-species temporal correlations, a non-parametric permutation test was performed. For each pair of human-pig components, the human time courses were fixed while the pig time courses were randomly circularly shifted (N=5000 permutations) and the correlations were recomputed at each iteration. This procedure generated a null distribution for each pair of components, against which the observed correlation was compared to obtain the p-values. The resulting p-values were corrected for multiple comparisons in the matrix using the FDR method (Benjamini-Hochberg) [79]. Hierarchical clustering was then performed using squared Euclidean distance and Ward’s method [80]. These analyses allowed us to characterize the organization of functional networks and investigate inter-species relationships.

To evaluate spatial coverage, non-noise components were converted into z-statistic images and thresholded using component-specific values derived from a mixture-model fit. These thresholds highlight voxels most strongly associated with each component and serve as a heuristic for separating structured signal from background noise, rather than representing formal voxel-wise statistical significance. The resulting thresholded maps were binarized and averaged across subjects to create probability maps.

Subsequently, voxel-wise probability maps from both pigs and humans were mapped onto standard brain surfaces (***SI, Supplementary Figure S4***). The probability maps for pigs were displayed on inflated cortical surface renderings of the Porcine Neurological Imaging Space atlas [31], while those for humans were visualized on inflated brain surface renderings of the Human Connectome Project Multi-modal atlas [52].

### Fiber Tractography Analysis in Pigs and Humans

Following preprocessing of diffusion-weighted imaging data in both species, diffusion tensors were estimated on a voxel-wise basis to characterize white matter microstructure. Whole-brain fiber tractography was performed using the fitted diffusion tensors and implemented using FSL [81] and MITK Diffusion [82]. Streamlines were generated with a deterministic tractography algorithm, applying step size and curvature thresholds optimized for each species. For pig data, fiber clustering was subsequently applied to the whole-brain tractogram to identify major fiber bundles and support tract-specific analyses. The resulting fiber pathways were then used to facilitate cross-species comparisons of whole-brain white matter architecture.

From the fitted tensors, voxel-wise maps of FA were derived, providing quantitative indices of tissue microstructural organization. At each voxel, the diffusion tensor is represented as a symmetric 3 *×* 3 matrix, which can be diagonalized to yield three eigenvalues: *λ*_1_, *λ*_2_, and *λ*_3_. Here, *λ*_1_ corresponds to the principal eigenvalue, while *λ*_2_ and *λ*_3_ correspond to the minor eigenvalues. FA quantifies the degree of diffusion anisotropy and is calculated as:

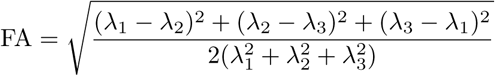

FA values range from 0, indicating completely isotropic diffusion, to 1, indicating highly anisotropic diffusion. These metrics provide complementary and biologically informative measures of white matter microstructural organization, facilitating comparisons across species.

### Estimation of Structural Connectivity (SC) in Pigs and Humans

SC matrices were constructed from diffusion-weighted MRI data using probabilistic tractography implemented in FSL’s PROBTRACKX [83]. Following diffusion preprocessing and diffusion model estimation for both species, as described previously [84, 85], probabilistic tractography was performed by initiating streamlines from every voxel within each atlas-defined seed region to estimate structural connectivity with all other brain regions while accounting for uncertainty in local fiber orientations.

For the humans, the cerebral cortex was parcellated using the 68-region Desikan-Killiany atlas [86]. For the pigs, brain regions were defined according to the selected 60-region Saikali pig brain atlas [76]. For each subject, the streamline counts between all pairs of atlas regions were assembled into a weighted SC matrix, resulting in a 68 × 68 matrix for humans and a 60 × 60 matrix for pigs. Because probabilistic tractography yields asymmetric connectivity estimates due to differences in seed and target sampling, each matrix was symmetrized by averaging the corresponding upper and lower triangular elements. Finally, each SC matrix was normalized by dividing every matrix element by the maximum connectivity value within that subject’s matrix, yielding normalized weighted connectivity values ranging from 0 to 1. These normalized SC matrices were used for all subsequent analyses. For visualization, the average matrices were log-transformed using log_10_(*SC* + 1).

### Cross-Species Comparison of SC in Pigs and Humans

To compare global SC patterns across species, subject-specific SC matrices were first normalized by network strength. For each subject, the SC matrix was divided by the mean nodal strength to minimize differences in overall connection magnitude between subjects. Species-specific group-average SC matrices were then obtained by averaging the normalized matrices across all subjects within each species. Because the human (68 regions) and pig (60 regions) atlases differed in the number of brain regions, direct edge-by-edge comparison was not performed. Instead, a SC fingerprint was generated for each subject by extracting the upper triangular elements of the normalized SC matrix (excluding the diagonal), corresponding to all unique pairwise connections. The edge weights were sorted in descending order to characterize the global distribution of structural connection strengths independent of anatomical correspondence.

Pairwise similarity between every pig and human subject was quantified by calculating the Pearson correlation coefficient between their ranked SC fingerprints, producing a 42 × 42 cross-species similarity matrix. To assess overall correspondence between species, the average SC fingerprints of pigs and humans were computed and compared using Spearman’s rank correlation. Statistical significance was evaluated using a two-sided test, and the resulting p-value was adjusted for multiple comparisons using the Benjamini-Hochberg FDR procedure. A density-colored scatter plot of the average ranked edge weights was generated using two-dimensional kernel density estimation to illustrate the correspondence between pig and human SC distributions.

## Data availability

The pig MRI data from this study are available upon reasonable request to the corresponding authors. The human GSP (https://www.neuroinfo.org/gsp) and HCP (https://www.humanconnectome.org/) datasets are publicly available.

## Code availability

The analyses in this study utilized the Group ICA of fMRI Toolbox (GIFT 4.0c, https://trendscenter.org/software/gift/) as well as additional software including MATLAB (R2020a, https://www.mathworks.com/), FSL (https://fsl.fmrib.ox.ac.uk/fsl/), ANTs (https://github.com/ANTsX/ANTs), AFNI (https://afni.nimh.nih.gov/), Connectome Workbench (https://github.com/Washington-University/workbench), MITK Diffusion (https://github.com/MIC-DKFZ/MITK-Diffusion), MRIcron (https://www.nitrc.org/projects/mricron) and SPM12 (https://www.fil.ion.ucl.ac.uk/spm/software/spm12/).

## Acknowledgments

This work was supported by the National Science Foundation (Grant 2112455), the National Institutes of Health (Grants R01MH123610 and R01MH119251 to V. D. Calhoun), and the National Institutes of Health, National Institute of Neurological Disorders and Stroke (Grants 1R21NS131526 and 1R21NS123732 to E. E. Kaiser., F. D. West and Q. Zhao).

## Contributions

Q. Li., V. D. Calhoun., Q. Zhao.: Conceptualization, Investigation, Software, Writing - Review & editing. I. Ahmed., M. F. Taber., M. H. Laballe., S. E. Sneed., E. E. Kaiser., F. D. West., Q. Zhao.: Pig MRI Data Acquisition and Preprocessing. Q. Li., I. Ahmed., M. F. Taber., M. H. Laballe., S. E. Sneed., E. E. Kaiser., F. D. West., V. D. Calhoun., Q. Zhao.: Writing - Review & editing. E. E. Kaiser., F. D. West., Q. Zhao., V. D. Calhoun.: Funding Acquisition.

## References

[1] Bart Ellenbroek and Jiun Youn. “Rodent models in neuroscience research: is it a rat race?” In: Disease models & mechanisms 9.10 (2016), pp. 1079–1087.

[2] Paul Manger, Jessica Cort, Naseem Ebrahim, Adelaya Goodman, Justine Henning, Mohamed Karolia, Stacey-Lee Rodrigues, and Goran Strkalj. “Is 21st century neuroscience too focussed on the rat/mouse model of brain function and dysfunction?” In: Frontiers in neuroanatomy 2 (2008), p. 329.

[3] Cheryl A Logan. “Commercial rodents in America: Standard animals, model animals, and biological diversity”. In: Brain, Behavior and Evolution 93.2–3 (2019), pp. 70–81.

[4] Judith R Homberg, Markus Wöhr, and Natalia Alenina. Comeback of the rat in biomedical research. 2017.

[5] Teresa Sierra-Arregui, Javier Llorente, Paula Giménez Minguez, Jan Tønnesen, and Olga Peñagarikano. “Neurobiological mechanisms of autism spectrum disorder and epilepsy, insights from animal models”. In: Neuroscience 445 (2020), pp. 69–82.

[6] Randall J Roper and Charles R Goodlett. “A new Down syndrome rat model races forward”. In: Trends in Genetics 38.11 (2022), pp. 1101–1102.

[7] Jürgen Götz, Liviu-Gabriel Bodea, and Michel Goedert. “Rodent models for Alzheimer disease”. In: Nature Reviews Neuroscience 19.10 (2018), pp. 583–598.

[8] Shubham Banga, Stefan D Heinze-Milne, and Susan E Howlett. “Rodent models of frailty and their application in preclinical research”. In: Mechanisms of ageing and development 179 (2019), pp. 1–10.

[9] IM Macrae. “Preclinical stroke research–advantages and disadvantages of the most common rodent models of focal ischaemia”. In: British journal of pharmacology 164.4 (2011), pp. 1062– 1078.

[10] Anand Gururajan, Andreas Reif, John F Cryan, and David A Slattery. “The future of rodent models in depression research”. In: Nature Reviews Neuroscience 20.11 (2019), pp. 686–701.

[11] EV Vykhovanets, MI Resnick, GT MacLennan, and S Gupta. “Experimental rodent models of prostatitis: limitations and potential”. In: Prostate cancer and prostatic diseases 10.1 (2007), pp. 15–29.

[12] Annabella Lear, Stuart N Baker, Hannah F Clarke, Angela C Roberts, Michael C Schmid, and Wendy Jarrett. “Understanding them to understand ourselves: The importance of NHP research for translational neuroscience”. In: Current Research in Neurobiology 3 (2022), p. 100049.

[13] Anna S Mitchell, Alexander Thiele, Christopher I Petkov, Angela Roberts, Trevor W Robbins, Wolfram Schultz, and Roger Lemon. “Continued need for non-human primate neuroscience research”. In: Current Biology 28.20 (2018), R1186–R1187.

[14] Pieter R Roelfsema and Stefan Treue. “Basic neuroscience research with nonhuman primates: a small but indispensable component of biomedical research”. In: Neuron 82.6 (2014), pp. 1200–1204.

[15] Alessandro Zanini, Audrey Dureux, Ravi S Menon, and Stefan Everling. “Mapping functional homologies between human and marmoset brain networks using movie-driven ultra-high field fMRI”. In: Communications Biology 9.1 (2026), p. 21.

[16] Qiang Li, Ting Xu, and Vince D Calhoun. “Functional correspondences in the human and marmoset visual cortex during movie watching: Insights from correlation, redundancy, and synergy”. In: Brain Research 1865 (2025), p. 149864.

[17] Qiang Li, Vince D Calhoun, and Armin Iraji. “Revealing complex functional topology brain network correspondences between humans and marmosets”. In: Neuroscience Letters 822 (2024), p. 137624.

[18] Dante Mantini, Uri Hasson, Viviana Betti, Mauro G Perrucci, Gian Luca Romani, Maurizio Corbetta, Guy A Orban, and Wim Vanduffel. “Interspecies activity correlations reveal functional correspondence between monkey and human brain areas”. In: Nature methods 9.3 (2012), pp. 277–282.

[19] Guoping Feng, Frances E Jensen, Henry T Greely, Hideyuki Okano, Stefan Treue, Angela C Roberts, James G Fox, Sarah Caddick, Mu-ming Poo, William T Newsome, et al. “Opportunities and limitations of genetically modified nonhuman primate models for neuroscience research”. In: Proceedings of the National Academy of Sciences 117.39 (2020), pp. 24022–24031.

[20] Michele A Basso, Aaron P Batista, Steve WC Chang, Katalin M Gothard, Cory T Miller, Karen J Parker, and Jan Zimmermann. “The future of nonhuman primate neuroscience: peril or possibilities?” In: Journal of Neuroscience 44.37 (2024).

[21] Yukitaka Tanaka, Hideaki Imai, Kenjiro Konno, Takaaki Miyagishima, Chisato Kubota, Sandra Puentes, Takeo Aoki, Hidekazu Hata, Kuniaki Takata, Yuhei Yoshimoto, et al. “Experimental model of lacunar infarction in the gyrencephalic brain of the miniature pig: neurological assessment and histological, immunohistochemical, and physiological evaluation of dynamic corticospinal tract deformation”. In: Stroke 39.1 (2008), pp. 205–212.

[22] Mitsunobu Nakamura, Hideaki Imai, Kenjiro Konno, Chisato Kubota, Koji Seki, Sandra Puentes, Ahmad Faried, Hideaki Yokoo, Hidekazu Hata, Yuhei Yoshimoto, et al. “Experimental investigation of encephalomyosynangiosis using gyrencephalic brain of the miniature pig: histopathological evaluation of dynamic reconstruction of vessels for functional anastomosis”. In: Journal of Neurosurgery: Pediatrics 3.6 (2009), pp. 488–495.

[23] David W Howells, Michelle J Porritt, Sarah SJ Rewell, Victoria O’collins, Emily S Sena, H Bart Van der Worp, Richard J Traystman, and Malcolm R Macleod. “Different strokes for different folks: the rich diversity of animal models of focal cerebral ischemia”. In: Journal of Cerebral Blood Flow & Metabolism 30.8 (2010), pp. 1412–1431.

[24] Holly A Kinder, Emily W Baker, and Franklin D West. “The pig as a preclinical traumatic brain injury model: current models, functional outcome measures, and translational detection strategies”. In: Neural regeneration research 14.3 (2019), pp. 413–424.

[25] Caijuan Li, Jun Li, Liangxue Lai, Shihua Li, and Sen Yan. “Genetically engineered pig models of neurological diseases”. In: Ageing and Neurodegenerative Diseases 2.3 (2022), N–A.

[26] Gregory Simchick, Alice Shen, Brandon Campbell, Hea Jin Park, Franklin D West, and Qun Zhao. “Pig brains have homologous resting-state networks with human brains”. In: Brain connectivity 9.7 (2019), pp. 566–579.

[27] Nina Reiter, Sarah Nistler, Lucas Hoffmann, Lars Bräuer, Friedrich Paulsen, and Silvia Budday. “Mechanical characterization of human versus porcine brain tissue under large strains”. In: Biomechanics and Modeling in Mechanobiology 25.2 (2026), p. 26.

[28] Morgan H LaBalle, Wenwu Sun, Ishfaque Ahmed, William D Reeves, Moira F Taber, Sydney E Sneed, Erin E Kaiser, Franklin D West, and Qun Zhao. “Longitudinal evaluation of white matter tracts post traumatic brain injury using a pediatric porcine model”. In: Journal of Neuroscience Methods (2026), p. 110684.

[29] Ishfaque Ahmed, William D Reeves, Morgan H Laballe, Moira F Taber, Sydney E Sneed, Erin E Kaiser, Franklin D West, and Qun Zhao. “A novel integration of brain structural and functional connectivity for identifying traumatic brain injury induced perturbations”. In: Journal of Neuroscience Methods (2025), p. 110459.

[30] Madison M Fagan, Kelly M Scheulin, Sydney E Sneed, Wenwu Sun, Christina B Welch, Savannah R Cheek, Erin E Kaiser, Qun Zhao, Kylee J Duberstein, and Franklin D West. “White Matter Integrity and Motor Function Disruption Due to Traumatic Brain Injury in Piglets: Impacts on Motor-Related Brain Fibers”. In: Brain sciences 14.3 (2024), p. 247.

[31] R Austin Benn, Rogier B Mars, Ting Xu, Jason R Yee, Luis Rodríguez-Esparragoza, Paula Montesinos, JP Manzano-Patron, Gonzalo Lopez-Martin, Valentin Fuster, Javier Sanchez-Gonzalez, et al. “Opening the pig to comparative neuroimaging: a common space approach contextualizes the pig and human structural connectome”. In: bioRxiv (2020), pp. 2020–10.

[32] Monika Baxa, Marian Hruska-Plochan, Stefan Juhas, Petr Vodicka, Antonin Pavlok, Jana Juhasova, Atsushi Miyanohara, Tetsuya Nejime, Jiri Klima, Monika Macakova, et al. “A transgenic minipig model of Huntington’s disease”. In: Journal of Huntington’s disease 2.1 (2013), pp. 47–68.

[33] Seung-Eun Lee, Hyuk Hyun, Mi-Ryung Park, Youngsok Choi, Yeo-Jin Son, Yun-Gwi Park, Sang-Gi Jeong, Min-Young Shin, Hee-Jin Ha, Hyun-Sok Hong, et al. “Production of transgenic pig as an Alzheimer’s disease model using a multi-cistronic vector system”. In: PLoS One 12.6 (2017), e0177933.

[34] MN Chieppa, A Perota, C Corona, A Grindatto, I Lagutina, E Vallino Costassa, Giovanna Lazzari, S Colleoni, R Duchi, Franco Lucchini, et al. “Modeling amyotrophic lateral sclerosis in hSOD1G93A transgenic swine”. In: Neurodegenerative diseases 13.4 (2014), pp. 246–254.

[35] Maria Shadrina and Petr Slominsky. “Modeling Parkinson’s disease: not only rodents?” In: Frontiers in aging neuroscience 13 (2021), p. 695718.

[36] Madison M Fagan, Christina B Welch, Kelly M Scheulin, Sydney E Sneed, Julie H Jeon, Morgane E Golan, Savannah R Cheek, Deborah A Barany, Georg Oeltzschner, Todd R Callaway, et al. “Fecal microbial transplantation limits neural injury severity and functional deficits in a pediatric piglet traumatic brain injury model”. In: Frontiers in Neuroscience 17 (2023), p. 1249539.

[37] Gregory Simchick, Kelly M Scheulin, Wenwu Sun, Sydney E Sneed, Madison M Fagan, Savannah R Cheek, Franklin D West, and Qun Zhao. “Detecting functional connectivity disruptions in a translational pediatric traumatic brain injury porcine model using resting-state and task-based fMRI”. In: Scientific Reports 11.1 (2021), p. 12406.

[38] Magdalena Hryhorowicz, Daniel Lipiński, Szymon Hryhorowicz, Agnieszka Nowak-Terpiłowska, Natalia Ryczek, and Joanna Zeyland. “Application of genetically engineered pigs in biomedical research”. In: Genes 11.6 (2020), p. 670.

[39] Gökhan Gün and Wilfried A Kues. “Current progress of genetically engineered pig models for biomedical research”. In: Bioresearch open access 3.6 (2014), pp. 255–264.

[40] Jialin Wei, Wen Zhang, Jie Li, Ye Jin, and Zhidong Qiu. “Application of the transgenic pig model in biomedical research: A review”. In: Frontiers in Cell and Developmental Biology 10 (2022), p. 1031812.

[41] Silja Zettler, Simone Renner, Elisabeth Kemter, Arne Hinrichs, Nikolai Klymiuk, Mattias Backman, Evamaria Olga Riedel, Christiane Mueller, Elisabeth Streckel, Christina Braun-Reichhart, et al. “A decade of experience with genetically tailored pig models for diabetes and metabolic research”. In: Animal reproduction 17.3 (2020), e20200064.

[42] Eugeni Roura, Sietse-Jan Koopmans, Jean-Paul Lallès, Isabelle Le Huerou-Luron, Nadia de Jager, Teun Schuurman, and David Val-Laillet. “Critical review evaluating the pig as a model for human nutritional physiology”. In: Nutrition research reviews 29.1 (2016), pp. 60–90.

[43] Hao Jia, Yuan Chang, and Jiangping Song. “The pig as an optimal animal model for cardiovascular research”. In: Lab Animal 53.6 (2024), pp. 136–147.

[44] George C Gabriel, William Devine, Bethany K Redel, Kristin M Whitworth, Melissa Samuel, Lee D Spate, Raissa F Cecil, Randall S Prather, Yijen Wu, Kevin D Wells, et al. “Cardiovascular development and congenital heart disease modeling in the pig”. In: Journal of the American Heart Association 10.14 (2021), e021631.

[45] Jean Noblet, Jean-Yves Dourmad, M Etienne, and Jean Le Dividich. “Energy metabolism in pregnant sows and newborn pigs”. In: Journal of Animal Science 75.10 (1997), pp. 2708–2714.

[46] Ishfaque Ahmed, William D Reeves, Wenwu Sun, Stephanie T Dubrof, Jillien G Zukaitis, Franklin D West, Hea Jin Park, and Qun Zhao. “Nutritional supplement induced modulations in the functional connectivity of a porcine brain”. In: Nutritional neuroscience 27.2 (2024), pp. 147–158.

[47] Muhammad M Mohiuddin, Bruno Reichart, Guerard W Byrne, and Christopher GA McGregor. “Current status of pig heart xenotransplantation”. In: International Journal of Surgery 23 (2015), pp. 234–239.

[48] Nader Moazami, Jeffrey M Stern, Karen Khalil, Jacqueline I Kim, Navneet Narula, Massimo Mangiola, Elaina P Weldon, Larisa Kagermazova, Les James, Nikki Lawson, et al. “Pig-to-human heart xenotransplantation in two recently deceased human recipients”. In: Nature medicine 29.8 (2023), pp. 1989–1997.

[49] David KC Cooper and Emanuele Cozzi. “Clinical pig heart xenotransplantation—where do we go from here?” In: Transplant International 37 (2024), p. 12592.

[50] Avram J Holmes, Marisa O Hollinshead, Timothy M O’keefe, Victor I Petrov, Gabriele R Fariello, Lawrence L Wald, Bruce Fischl, Bruce R Rosen, Ross W Mair, Joshua L Roffman, et al. “Brain Genomics Superstruct Project initial data release with structural, functional, and behavioral measures”. In: Scientific data 2.1 (2015), pp. 1–16.

[51] David C Van Essen, Stephen M Smith, Deanna M Barch, Timothy EJ Behrens, Essa Yacoub, Kamil Ugurbil, Wu-Minn HCP Consortium, et al. “The WU-Minn human connectome project: an overview”. In: Neuroimage 80 (2013), pp. 62–79.

[52] Matthew F Glasser, Timothy S Coalson, Emma C Robinson, Carl D Hacker, John Harwell, Essa Yacoub, Kamil Ugurbil, Jesper Andersson, Christian F Beckmann, Mark Jenkinson, et al. “A multi-modal parcellation of human cerebral cortex”. In: Nature 536.7615 (2016), pp. 171–178.

[53] Joan K Lunney, Angelica Van Goor, Kristen E Walker, Taylor Hailstock, Jasmine Franklin, and Chaohui Dai. “Importance of the pig as a human biomedical model”. In: Science translational medicine 13.621 (2021), eabd5758.

[54] Nanna Marie Lind, Anette Moustgaard, Jacob Jelsing, Gabor Vajta, Paul Cumming, and Axel K Hansen. “The use of pigs in neuroscience: modeling brain disorders”. In: Neuroscience & Biobehavioral Reviews 31.5 (2007), pp. 728–751.

[55] Wenwu Sun, Ishfaque Ahmed, Stephanie T Dubrof, Hea Jin Park, Franklin D West, and Qun Zhao. “Affinity of structural white matter tracts between infant and adult pig”. In: Journal of Neuroscience Methods 406 (2024), p. 110134.

[56] Elise T Gieling, Teun Schuurman, Rebecca E Nordquist, and F Josef van der Staay. “The pig as a model animal for studying cognition and neurobehavioral disorders”. In: Molecular and functional models in neuropsychiatry (2011), pp. 359–383.

[57] Alesa H Netzley and Galit Pelled. “The pig as a translational animal model for biobehavioral and neurotrauma research”. In: Biomedicines 11.8 (2023), p. 2165.

[58] Qiang Li, Greg Ver Steeg, Shujian Yu, and Jesus Malo. “Functional connectome of the human brain with total correlation”. In: Entropy 24.12 (2022), p. 1725.

[59] Qiang Li, Greg Ver Steeg, and Jesús Malo. “Functional connectivity via total correlation: Analytical results in visual areas”. In: Neurocomputing 571 (2024), p. 127143.

[60] Qiang Li, Jingyu Liu, Godfrey D Pearlson, Jiayu Chen, Yu-Ping Wang, Jessica A Turner, and Vince D Calhoun. “Spatiotemporal complexity in the psychotic brain”. In: Molecular Psychiatry (2025), pp. 1–15.

[61] Qiang Li, Shujian Yu, Jesus Malo, Godfrey D Pearlson, Yu-Ping Wang, and Vince D Calhoun. “Higher-Order Triadic Interactions: Insights Into the Multiscale Network Organization in Schizophrenia”. In: Human Brain Mapping 46.16 (2025), e70399.

[62] R Matthew Hutchison, Thilo Womelsdorf, Elena A Allen, Peter A Bandettini, Vince D Calhoun, Maurizio Corbetta, Stefania Della Penna, Jeff H Duyn, Gary H Glover, Javier Gonzalez-Castillo, et al. “Dynamic functional connectivity: promise, issues, and interpretations”. In: Neuroimage 80 (2013), pp. 360–378.

[63] Michelle M McCarthy, ShiNung Ching, Miles A Whittington, and Nancy Kopell. “Dynamical changes in neurological diseases and anesthesia”. In: Current opinion in neurobiology 22.4 (2012), pp. 693–703.

[64] UnCheol Lee and George A Mashour. “The role of network science in the study of anesthetic state transitions”. In: Anesthesiology 129.5 (2018), p. 1029.

[65] Mohsen Bahrami, Paul J Laurienti, Heather M Shappell, and Sean L Simpson. “Brain network analysis: A review on multivariate analytical methods”. In: Brain connectivity 13.2 (2023), pp. 64–79.

[66] Christopher J Honey, Jean-Philippe Thivierge, and Olaf Sporns. “Can structure predict function in the human brain?” In: Neuroimage 52.3 (2010), pp. 766–776.

[67] Arsène Ella, David A Barrière, Hans Adriaensen, David N Palmer, Tracy R Melzer, Nadia L Mitchell, and Matthieu Keller. “The development of brain magnetic resonance approaches in large animal models for preclinical research”. In: Animal Frontiers 9.3 (2019), pp. 44–51.

[68] Aage Kristian Olsen Alstrup and Michael Winterdahl. “Imaging techniques in large animals”. In: Scandinavian Journal of Laboratory Animal Science 36.1 (2009), pp. 55–66.

[69] Hui Zhang, Torben Schneider, Claudia A Wheeler-Kingshott, and Daniel C Alexander. “NODDI: practical in vivo neurite orientation dispersion and density imaging of the human brain”. In: Neuroimage 61.4 (2012), pp. 1000–1016.

[70] Eduardo Caverzasi, Nico Papinutto, Antonella Castellano, Alyssa H Zhu, Paola Scifo, Marco Riva, Lorenzo Bello, Andrea Falini, Aditya Bharatha, and Roland G Henry. “Neurite orientation dispersion and density imaging color maps to characterize brain diffusion in neurologic disorders”. In: Journal of Neuroimaging 26.5 (2016), pp. 494–498.

[71] Vince D Calhoun, Tülay Adalı, Godfrey D Pearlson, and James J Pekar. “A Method for Making Group Inferences from Functional MRI Data Using Independent Component Analysis”. In: Human Brain Mapping 14 (2001), pp. 140–151.

[72] Srinivas Rachakonda, Rogers F Silva, Jingyu Liu, and Vince D Calhoun. “Memory efficient PCA methods for large group ICA”. In: Frontiers in neuroscience 10 (2016), p. 17.

[73] Erik Barry Erhardt, Srinivas Rachakonda, Edward J Bedrick, Elena A Allen, Tülay Adali, and Vince D Calhoun. “Comparison of multi-subject ICA methods for analysis of fMRI data”. In: Human brain mapping 32.12 (2011), pp. 2075–2095.

[74] Anthony J Bell and Terrence J Sejnowski. “An information-maximization approach to blind separation and blind deconvolution”. In: Neural computation 7.6 (1995), pp. 1129–1159.

[75] Johan Himberg, Aapo Hyvärinen, and Fabrizio Esposito. “Validating the independent components of neuroimaging time series via clustering and visualization”. In: Neuroimage 22.3 (2004), pp. 1214–1222.

[76] Stéphan Saikali, Paul Meurice, Paul Sauleau, Pierre-Antoine Eliat, Pascale Bellaud, Gwenaelle Randuineau, Marc Vérin, and Charles-Henri Malbert. “A three-dimensional digital segmented and deformable brain atlas of the domestic pig”. In: Journal of neuroscience methods 192.1 (2010), pp. 102–109.

[77] Stephen M Smith, Peter T Fox, Karla L Miller, David C Glahn, P Mickle Fox, Clare E Mackay, Nicola Filippini, Kate E Watkins, Roberto Toro, Angela R Laird, et al. “Correspondence of the brain’s functional architecture during activation and rest”. In: Proceedings of the national academy of sciences 106.31 (2009), pp. 13040–13045.

[78] and others. In: Human brain mapping 44.17 (2023), pp. 5729–5748.

[79] David Thissen, Lynne Steinberg, and Daniel Kuang. “Quick and easy implementation of the Benjamini-Hochberg procedure for controlling the false positive rate in multiple comparisons”. In: Journal of educational and behavioral statistics 27.1 (2002), pp. 77–83.

[80] Pedro Contreras and Fionn Murtagh. “Hierarchical clustering”. In: Handbook of cluster analysis (2015), pp. 103–123.

[81] Mark Jenkinson, Christian F Beckmann, Timothy EJ Behrens, Mark W Woolrich, and Stephen M Smith. “Fsl”. In: Neuroimage 62.2 (2012), pp. 782–790.

[82] Peter F Neher, Bram Stieltjes, Marco Reisert, Ignaz Reicht, Hans-Peter Meinzer, and Klaus H Fritzsche. “MITK global tractography”. In: Medical Imaging 2012: Image Processing. Vol. 8314. SPIE. 2012, pp. 1232–1237.

[83] Stephen M Smith, Mark Jenkinson, Mark W Woolrich, Christian F Beckmann, Timothy EJ Behrens, Heidi Johansen-Berg, Peter R Bannister, Marilena De Luca, Ivana Drobnjak, David E Flitney, et al. “Advances in functional and structural MR image analysis and implementation as FSL”. In: Neuroimage 23 (2004), S208–S219.

[84] Ishfaque Ahmed, Morgan H LaBalle, Moira F Taber, Sydney E Sneed, Erin E Kaiser, Franklin D West, Taotao Wu, and Qun Zhao. “Assessment of Coupled Phase Oscillators-Based Modeling in Swine Brain Connectome”. In: Journal of Neuroscience Methods (2026), p. 110811.

[85] John Fallon, Phillip GD Ward, Linden Parkes, Stuart Oldham, Aurina Arnatkevičiūte, Alex Fornito, and Ben D Fulcher. “Timescales of spontaneous fMRI fluctuations relate to structural connectivity in the brain”. In: Network neuroscience 4.3 (2020), pp. 788–806.

[86] Rahul S Desikan, Florent Ségonne, Bruce Fischl, Brian T Quinn, Bradford C Dickerson, Deborah Blacker, Randy L Buckner, Anders M Dale, R Paul Maguire, Bradley T Hyman, et al. “An automated labeling system for subdividing the human cerebral cortex on MRI scans into gyral based regions of interest”. In: Neuroimage 31.3 (2006), pp. 968–980.

